# A CLN6-CRMP2-KLC4 complex regulates anterograde ER-derived vesicle trafficking in cortical neurites

**DOI:** 10.1101/2021.09.16.460653

**Authors:** SY Koh, JT Cain, H Magee, KA White, M Rechtzigel, B Meyerink, H Leppert, DJ Timm, JP Morgan, TB Johnson, B Grove, R Khanna, K Hensley, J Brudvig, JM Weimer

## Abstract

As neurons establish extensive connections throughout the central nervous system, the transport of cargo along the microtubule network of the axon is crucial for differentiation and homeostasis. Specifically, building blocks such as membrane and cytoskeletal components, organelles, transmembrane receptors, adhesion molecules, and peptide neurotransmitters all require proper transport to the presynaptic compartment. Here, we identify a novel complex regulating vesicular endoplasmic reticulum transport in neurites, composed of <u>C</u>LN6: an ER-associated protein of relatively unknown function implicated in CLN6-Batten disease; <u>C</u>RMP2: a tubulin binding protein important in regulating neurite microtubule dynamics; and <u>K</u>LC4: a classic transport motor protein. We show that this “CCK” complex allows ER-derived vesicles to migrate to the distal end of the axon, aiding in proper neurite outgrowth and arborization. In the absence of CLN6, the CCK complex does not function effectively, leading to reduced vesicular transport, stunted neurite outgrowth, and deficits in CRMP2 binding to other protein partners. Treatment with a CRMP2 modulating compound, lanthionine ketimine ester, partially restores these deficits in CLN6-deficient mouse neurons, indicating that stabilization of CRMP2 interacting partners may prove beneficial in lieu of complete restoration of the CCK complex. Taken together, these findings reveal a novel mechanism of ER-derived vesicle transport in the axon and provide new insights into therapeutic targets for neurodegenerative disease.

## INTRODUCTION

Neurons relay signals across vast distances using electrical impulses transmitted along axons. A consequence of these long cellular processes is that neurons require specialized forms of transport to deliver cargoes to these distal sites [1]. During neuronal differentiation, transport systems supply growing neurites with structural building blocks such as membranes [2], organelles including lysosomes and mitochondria, and a plethora of transmembrane receptors and adhesion molecules which facilitate proper establishment of the neuronal arbor [3, 4]. Once neurons have established these distal appendages, transport is similarly required for synaptic homeostasis, plastic remodeling, and for the relay of intracellular signals [1].

Within axons, transport of both cytoplasmic and vesicle-associated cargoes is accomplished through microtubule-mediated transport. Anterograde axonal transport, which carries a large variety of newly synthesized cargoes including cytosolic and vesicular proteins, organelles, and RNAs from the cell body to distal sites including the axon terminal, is mediated by kinesins. Kinesins are heterotetrameric transport motors consisting of two identical kinesin heavy chain motors, along with variable kinesin light chain adapter proteins that couple the motors to various cargoes (KLC1-4) [5, 6]. While several specific examples of KLC-associated transport complexes have been identified, including some present on ER-derived vesicles, it remains unknown how most cargoes, particularly those in vesicles, link to the KLC components of kinesin motors [7-9]. Likewise, while it has been shown that various KLCs are critical for the establishment of neuronal polarity and axon outgrowth, many questions remain regarding what cargoes and adapter proteins are involved in these roles [10-13].

Here, we shed light on this question by identifying a novel complex composed of KLC4, CRMP2, and CLN6. CRMP2 (collapsin response mediator protein-2, also known as DPYSL2) has emerged as an important mechanistic link between microtubule transport and neurite outgrowth and branching [14-16]. CRMP2 associates with KLC1 to bind kinesin motors to the SRA1/WAVE complex, which is critical for actin dynamics in developing axons [17]. Similarly, tubulin is transported within axons through a CRMP2/KLC1 complex, demonstrating that CRMP2 facilitates the transport of diverse cargoes [18]. CRMP2 has also been shown to associate with a variety of axonal proteins, including transmembrane vesicular proteins, but the functional relevance of these interactions is unknown [19, 20].

Additionally, CRMP2 interacts with the transmembrane ER-associated protein CLN6 [21, 22]. CLN6 is ubiquitously expressed and its loss of function results in CLN6-Batten disease, a lysosmal storage disorder characterized by neurodegeneration and the accumulation of ceroid/lipofuscin material throughout cells of various organs and tissues. This autosomal recessive disease manifests in early childhood with broad neurological symptoms including motor disturbance, vision impairment, and developmental regression, typically leading to death by the second decade of life [23-26]. Recently, CLN6 has been shown to work cooperatively with CLN8, the protein product of the causative gene implicated in CLN8-Batten Disease, in ER-to-Golgi transport of lysosomal enzymes [27]. While this role was discovered in non-neuronal cells and lysosomal dysfunction caused by loss of either CLN6 or CLN8 is predicted to affect all cell types, the specific vulnerability of neuronal populations in Batten disease suggests that these proteins may have undescribed neuron-specific functions.

Here, we show that CLN6 interacts with a CRMP2-KLC4 complex to regulate anterograde axonal transport in developing neurons. In the absence of CLN6, CRMP2 is unable to properly associate with key neuronal protein partners, disrupting axonal transport and axonal arborization. We show that treatment with a CRMP2 modulator, lanthionine ketimine ester (LKE), is able to rescue some, but not all, of these deficits in primary cultured neurons and an animal model with CLN6 deficiency. This work identifies the CCK complex as a critical regulator of axonal transport and neurite morphogenesis, identifies a novel, neuron-specific function of CLN6 that has direct implications in disease pathogenesis, and suggests potential avenues for future therapeutics in Batten disease and other neurodegenerative disorders.

## RESULTS

### CLN6 interacts with CRMP2 and KLC4 to regulate axonal transport of ER-derived vesicles

Mutations in *CLN6* lead to a host of neurological and cognitive deficits early in life, suggesting that developmental processes could be disrupted. Since the known molecular partners and functions of CLN6 cannot explain the neuron-centric phenotypes that result from CLN6 deficiency, we sought to identify novel CLN6 interactors with roles in neurodevelopment. We performed a yeast two-hybrid screen with the N-terminal cytosolic domain of CLN6 as a bait protein, and identified a putative interaction with KLC4, a kinesin light-chain protein which participates in microtubule transport (**Figure 1a**). Since CLN6 has also been shown to interact with CRMP2, which interacts with KLC proteins and regulates the transport of tubulin and the SRA/WAVE complex, we asked whether these three proteins could be participating in a complex to regulate axonal transport of ER-derived vesicles [9, 17]. As we could not validate any commercial antibodies reported to target CLN6, we utilized the expression of a CLN6-myc construct to validate candidate interactions. When CLN6-myc was expressed in PC12 cells, KLC4 co-immunoprecipitated with both CLN6-myc and CRMP2 (**Figure 1b**). Likewise, when we examined the localization patterns of CLN6-myc, CRMP2, and KLC4 in cultured primary cortical neurons, we identified numerous axonal vesicles which were positive for all three proteins (**Figure 1c**). We refer to this <u>C</u>LN6-<u>C</u>RMP2-<u>K</u>LC4 complex as the CCK complex.

**Figure 1.**
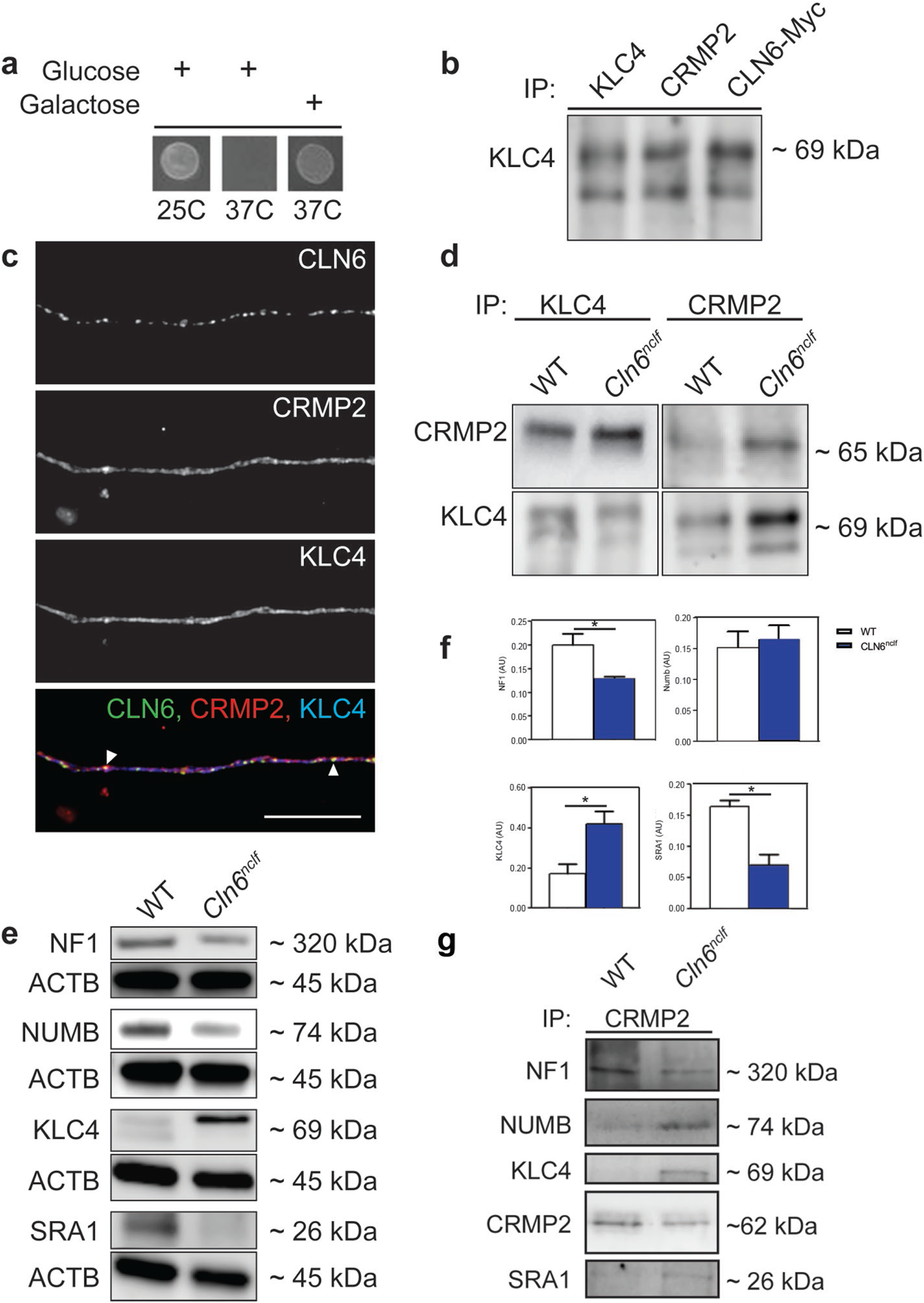
CLN6, CRMP2, and KLC4 complex together. Demonstration of yeast two hybrid screening with CLN6 fused to hSos screened against a myristoylated KLC4. In the presence of galactose, colony growth is restored at 37C, indicating an interaction between CLN6 and KLC4. b) CRMP2 and CLN6-Myc immunoprecipitate with the motor protein KLC4. c) CLN6, CRMP2, and KLC4 colocalize in the axon of mouse cortical neurons (white arrows). Scale bar 10μm. d) Loss of CLN6 results in enhanced CRMP2-KLC4 binding, as measured by co-immunoprecipitation. e) Loss of CLN6 results in altered total protein levels of various CRMP2 interacting proteins, as measured by Western blot. f) Quantification of e. g) Loss of CLN6 results in altered CRMP2 binding dynamics as measured by co-immunoprecipitation, distinct from total protein levels changes of various protein partners. Mean ± SEM, unpaired student’s t-test. *p<0.05.

To interrogate functional roles for CLN6 and the CCK complex, we utilized a naturally occurring mouse model that harbors an insertion mutation in exon 4 of *Cln6* resulting in a frameshift and premature stop codon, mirroring a common mutation found in human CLN6-Batten disease patients [24, 28]. This mutation (*Cln6*^*nclf*^) results in nonsense mediated decay-targeted degradation of the mutant transcript and undetectable levels of protein, making the model a powerful tool for examining the consequences of CLN6 deficiency [29, 30]. We performed co-immunoprecipitations on adult mouse brain lysates from wild type and *Cln6*^*nclf*^ mice to examine the CRMP2-KLC4 interaction. Surprisingly, we detected greater levels of CRMP2-KLC4 interaction when CLN6 was absent, suggesting that CLN6 may negatively regulate coupling of CRMP2 to KLC4 and the microtubule network (**Figure 1d**). We also asked whether CLN6 deficiency altered the expression or binding patterns of other CRMP2 partners, since CRMP2 interactors have been shown to be perturbed in various disease states [17, 19, 20]. *Cln6*^*nclf*^ mice had decreased total levels of NF1 and SRA1 and increased levels of KLC4 (**Figure 1e-f**). Interestingly, while interactions between CRMP2, NF1, and KLC4 were in line with changes in total protein expression in the absence of CLN6, interactions between CRMP2, NUMB, and SRA1 were increased, indicating that loss of CLN6 may change CRMP2’s ability to bind to target proteins or cargo (**Figure 1g**). As multiple phosphorylation sites on CRMP2 have been shown to modulate interactions with substrates, we examined whether CRMP2’s phosphorylation status was modulated in the absence of CLN6. Total levels of CRMP2 and pCRMP2 at multiple residues were unchanged, suggesting that CLN6 influences CRMP2 interactions without altering CRMP2 phosphorylation patterns (**Supp. Fig. 1a-b**).

CRMP2 and KLC4 have well established functions in axonal transport, but roles for CLN6 in this process have not been described. Since CLN6 is a resident ER transmembrane protein involved in ER to Golgi vesicle transport of lysosomal enzymes, we asked whether the CCK complex could be regulating the axonal transport of ER-derived vesicles [9, 27, 31, 32]. Mouse cortical neurons were cultured and labeled for ER-derived vesicles in live cells with ER-tracker. *Cln6*^*nclf*^ neurons had significantly fewer ER-derived vesicles within axons and an accumulation of ER-tracker signal in the neuronal soma, indicating a dysfunction in transporting ER-derived vesicles out of the soma (**Figure 2a-e**). Importantly, ER-derived vesicles from *Cln6*^*nclf*^ cells that did move along the axon were transported away from the soma (anterograde) at similar rates to wild type counterparts (**Figure 2f-h)**. As CLN6-Batten disease is a lysosomal storage disorder, we asked whether CLN6 deficiency also led to altered lysosome trafficking. Mouse cortical neurons were cultured and labeled for lysosomes with Lamp1 and monitored for velocity along the axon using LysoTracker. Interestingly, CLN6 deficiency caused no changes in either lysosome number or velocity along axons (**Figure 2i-j**), indicating that deficits are specific to another class of vesicles in *Cln6*^*nclf*^ cells.

**Figure 2.**
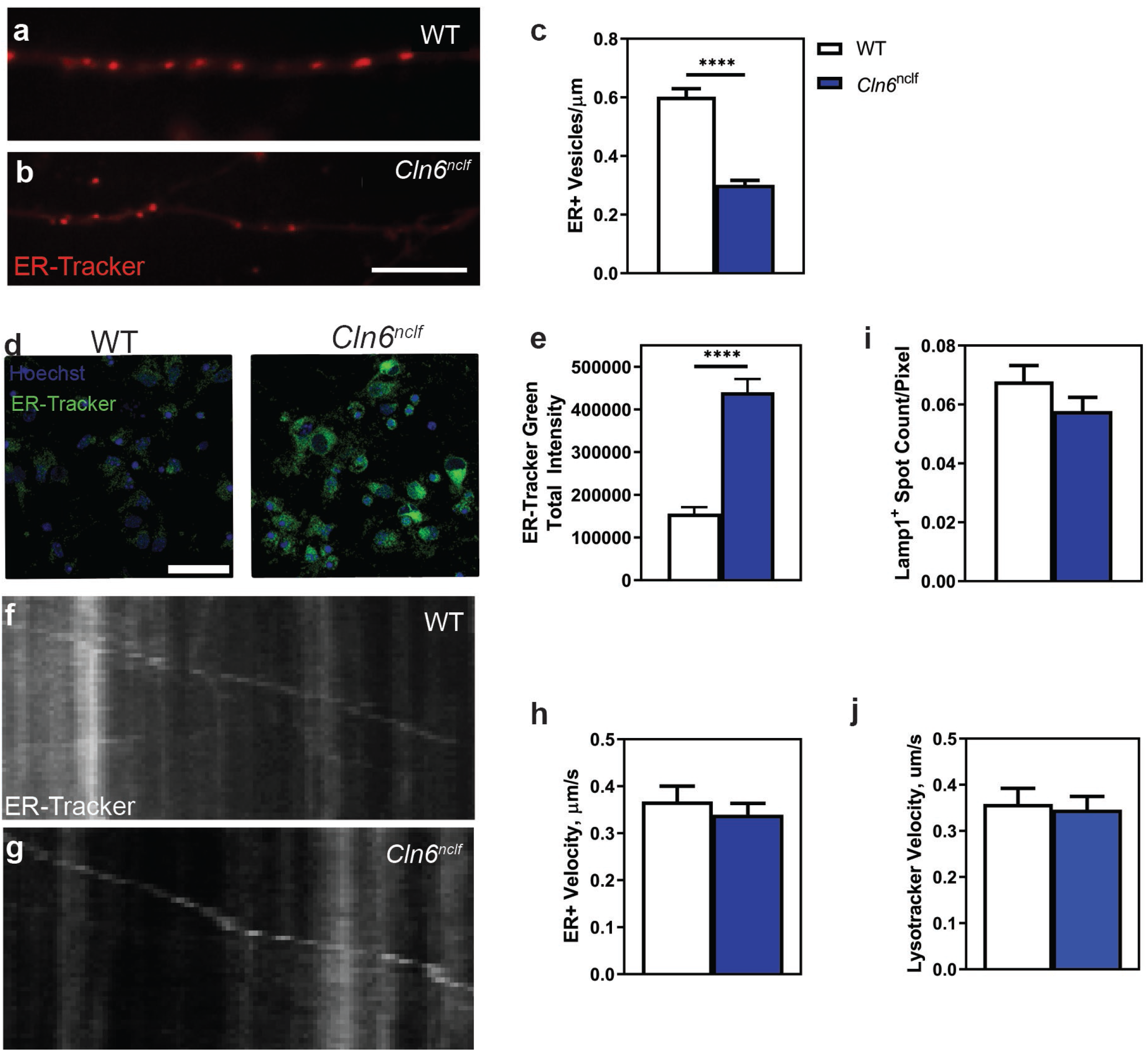
CLN6 deficiency leads to altered ER vesicle transport dynamics. a-c) Loss of CLN6 results in fewer ER positive vesicles in the axon of mouse cortical neurons, as monitored via ER-tracker. d-e) Loss of CLN6 results in the accumulation of ER-positive vesicles in the soma. f-h) Loss of CLN6 does not change ER positive vesicle velocity in the axon of mouse cortical neurons. g) Loss of CLN6 does not alter Lamp1^+^ vesicle counts in the axon of mouse cortical neurons. h) Loss of CLN6 does not alter lysosomal vesicle velocities in the axon of mouse cortical neurons, as measured by LysoTracker. Mean ±SEM, unpaired student’s t-test for all graphs. ****p<0.0001. Scale bar for images a-b: 5μm, d: 50 μm.

As ER-derived vesicles carry a variety of cargo, we asked whether the trafficking of lysosomal enzymes could be disrupted in the absence of CLN6. Synaptosomes from adult wild type and *Cln6*^*nclf*^ brains were isolated and measured for lysosomal enzyme activity, focusing on enzymes altered in other forms of Batten disease. While Cathepsin D (CTSD) activity was only slightly decreased in *Cln6*^*nclf*^ whole brain extracts, *Cln6*^*nclf*^ synaptosomes had approximately only 50% of CTSD activity as compared to wild type (**Figure 3a-b**). Similarly, palmitoyl-protein thioesterase 1 (PPT1) activity was not significantly decreased in *Cln6*^*nclf*^ whole brain extracts but was severely decreased in *Cln6*^*nclf*^ synaptosomes by approximately 75% (**Figure 3c-d**). While we did not identify significant changes in tripeptidyl-peptidase 1 (TPP1) activity, whole brain extracts trended towards higher levels in *Cln6*^*nclf*^ samples (p=0.0583) and lower levels in *Cln6*^*nclf*^ synaptosomes (p=0.0901), indicating a common theme of reduced lysosomal enzyme activity in neuronal processes of *Cln6* deficient animals (**Figure 3e-f**). Together, these results demonstrate that, in the absence of CLN6, the activity of a variety of lysosomal enzymes is compromised specifically in the synaptic compartment, suggesting that the trafficking role of CLN6 may be necessary for the transport of lysosomal enzymes to distal sites within neurites, perhaps carried as cargo in ER-derived vesicles.

**Figure 3.**
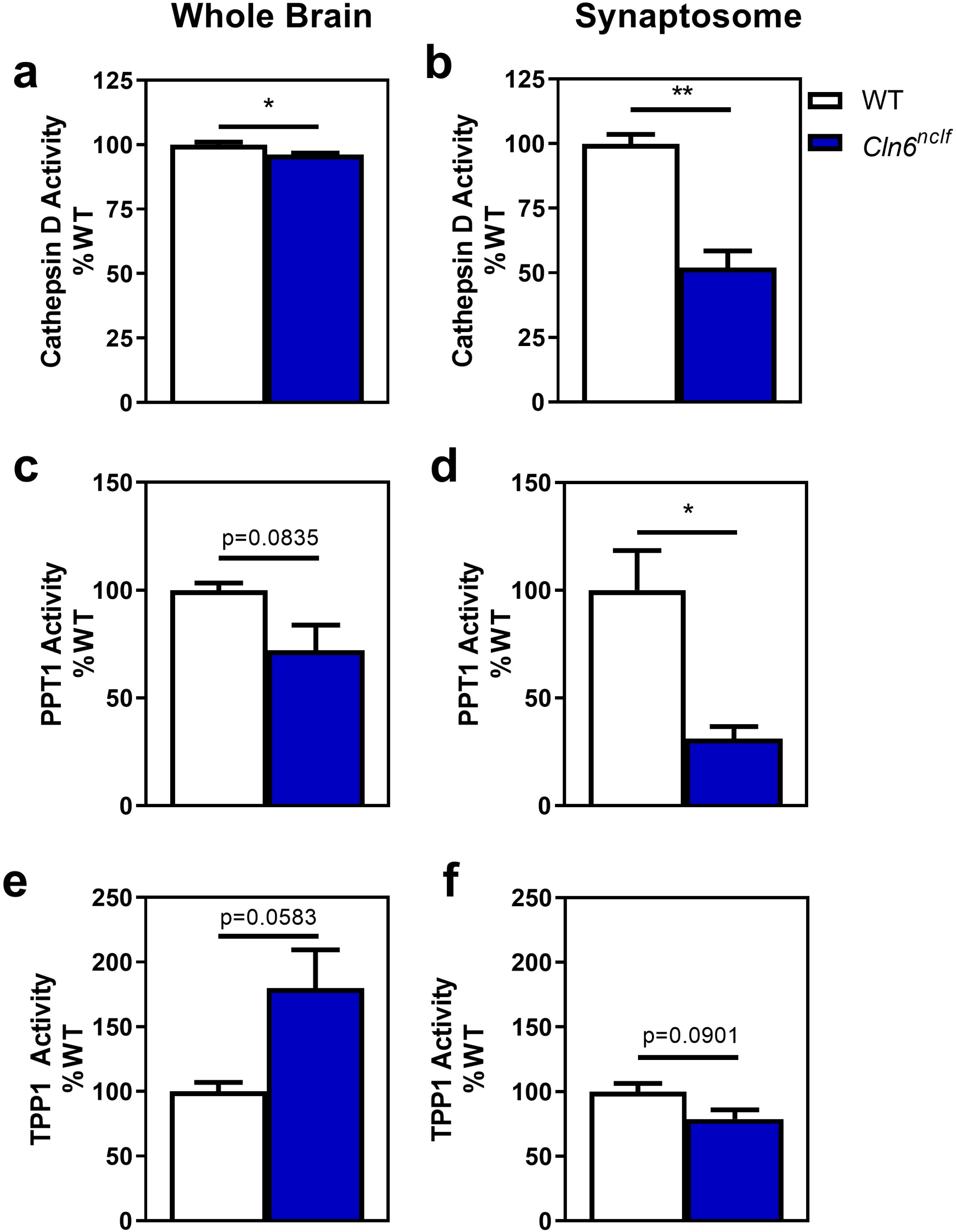
CLN6 deficiency is necessary for proper lysosomal enzyme trafficking. a) Loss of CLN6 leads to lower levels of Cathepsin D enzyme activity in both the whole brain and synaptosome. b) Loss of CLN6 leads to lower levels of PPT1 in the synaptosome, though not in the whole brain. c) Loss of CLN6 does not significantly alter levels of TPP1 enzyme in either the brain or synaptosome. Mean ±SEM, unpaired student’s t-test for all graphs. *p<0.05, **p<0.01.

### CLN6 is required for neurite morphogenesis

Throughout development, axonal transport of the proper cargo is critical for neuronal polarization as well as neurite extension and branching [29, 31, 33-35]. Transport machinery mediates the partitioning of axon- and dendrite-specific cargoes which supply growing neurites with structural components required for extension. Since the CLN6 regulates axonal transport, we asked whether neurite initiation and morphogenesis are compromised in the absence of CLN6. Primary cortical neurons from wild type and *Cln6*^*nclf*^ mice, electroporated with pCAG-GFP, were cultured and examined for initial neurite polarization and extension in GFP^+^ cells. When cultured at low density (25,000 cells/well), *Cln6*^*nclf*^ neurons exhibited delayed polarization and maturation, as evidenced by the lack of a defined axon and short, immature neurites (**Figure 4a-f**). By 7 days *in vitro* (7 DIV), this deficiency was qualitatively rectified (**Figure 4g-h**). However, when cultured for 7 DIV at higher densities (50,000 cells/well) that favor more robust growth, *Cln6*^*nclf*^ neurons had significantly shorter axons and dendrites, significantly less branching of axons and dendrites, and simplified arborization patterns when analyzed via Sholl analysis (**Figure 4i-q**).

**Figure 4.**
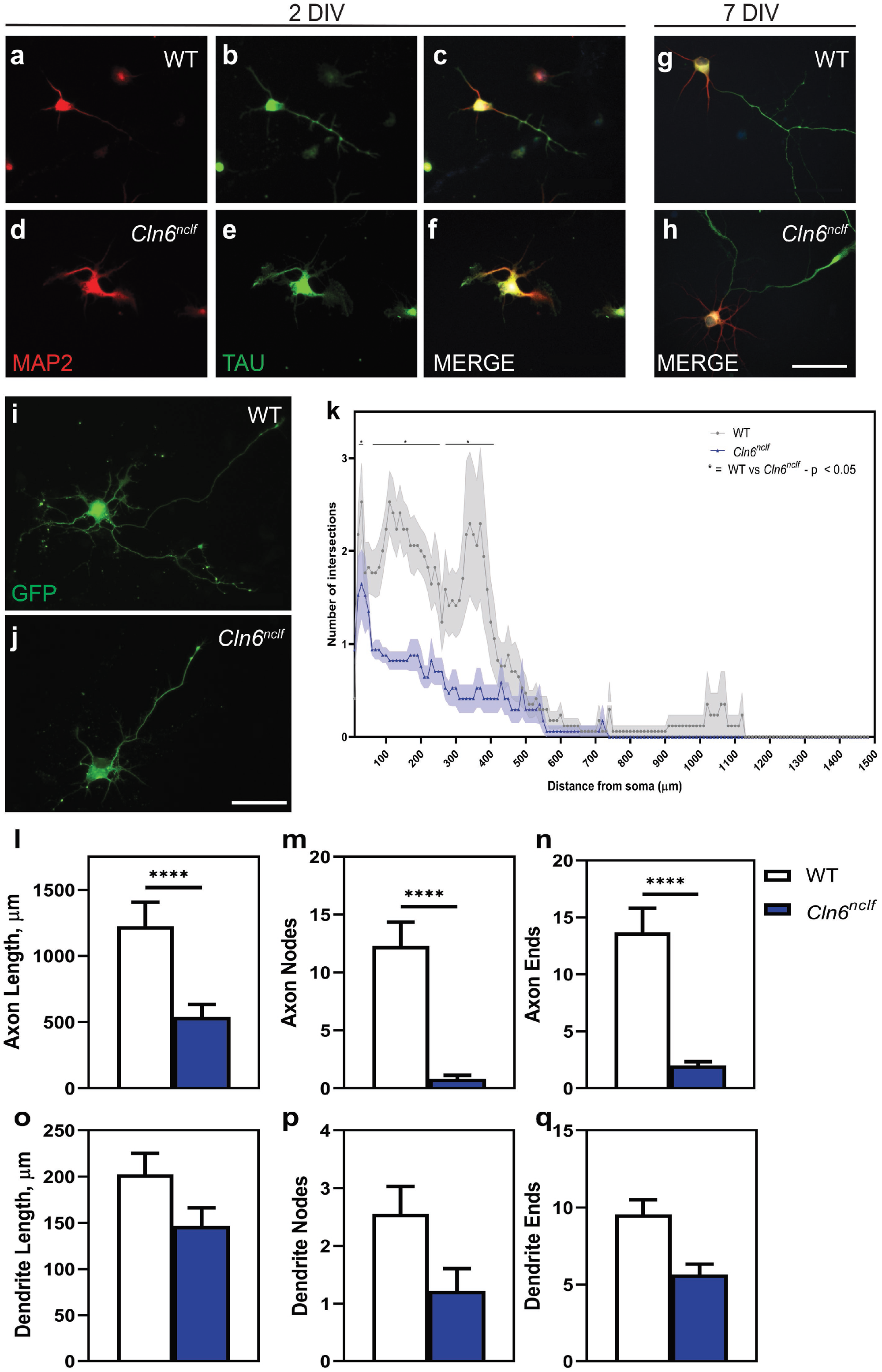
CLN6 is required for neurite outgrowth and establishment of neurite arborization. a-f) *Cln6*^*nclf*^ neurons have deficits in neurite outgrowth at 2 days *in vitro* (DIV) when plated at low density (25,000 cells/well). g,h) *Cln6*^*nclf*^ neurite deficits are restored by 7 DIV. i-j) *Cln6*^*nclf*^ neurons plated at a higher density (50,000 cells/well) continue to have neurite outgrowth deficits at 7 DIV. k) Sholl analysis of 7 DIV neurons plated at 50,000 cells/well, showing deficits in neurite arborization of *Cln6*^*nclf*^ neurons. l-q) Deficits in a variety of axon and dendrite parameters in *Cln6*^*nclf*^ neurons, particularly in axon length, number of axon nodes, and number of axon ends. Mean ± SEM. Two-way ANOVA with Bonferroni’s post hoc test for Sholl Analysis; unpaired student’s t-test for l-q. *p<0.05, ****p<0.0001. Scale bars 50μm.

To investigate *in vivo* axon phenotypes in the absence of CLN6, we used transmission electron microscopy to examine corpus callosum ultrastructure in sagittal brain sections, which capture cross sections of callosal axons as they cross the midline. At 1 month of age, axon density within the corpus callosum was not changed in *Cln6*^*nclf*^ brains; however, the average myelin sheath thickness was significantly but mildly decreased (**Figure 5a-b**). At 10 months of age, when more axons have crossed the midline, both axon caliber and myelin sheath thickness were significantly decreased in *Cln6*^*nclf*^ brains, as was overall volume of the corpus callosum (**Figure 5 c-d**). The decrease in corpus callosum volume and myelin corroborate changes seen with magnetic resonance imaging and diffusion tensor imaging of *Cln6*^*nclf*^ mice [36]. Taken together, these *in vitro* and *in vivo* results suggest that axon extension and branching are compromised in the absence of Cln6 and that these cellular deficiencies lead to defects in corpus callosum architecture. While we did not explore the myelination phenotype in greater detail, our results also suggest that *Cln6*^*nclf*^ oligodendrocytes may have defects in process extension, manifesting as reduced myelin sheath thickness in the corpus callosum. This would agree with a role for the CCK complex in microtubule transport since oligodendrocytes rely heavily on microtubule-dependent transport of mRNAs to distal sites in the myelin sheath [34].

**Figure 5.**
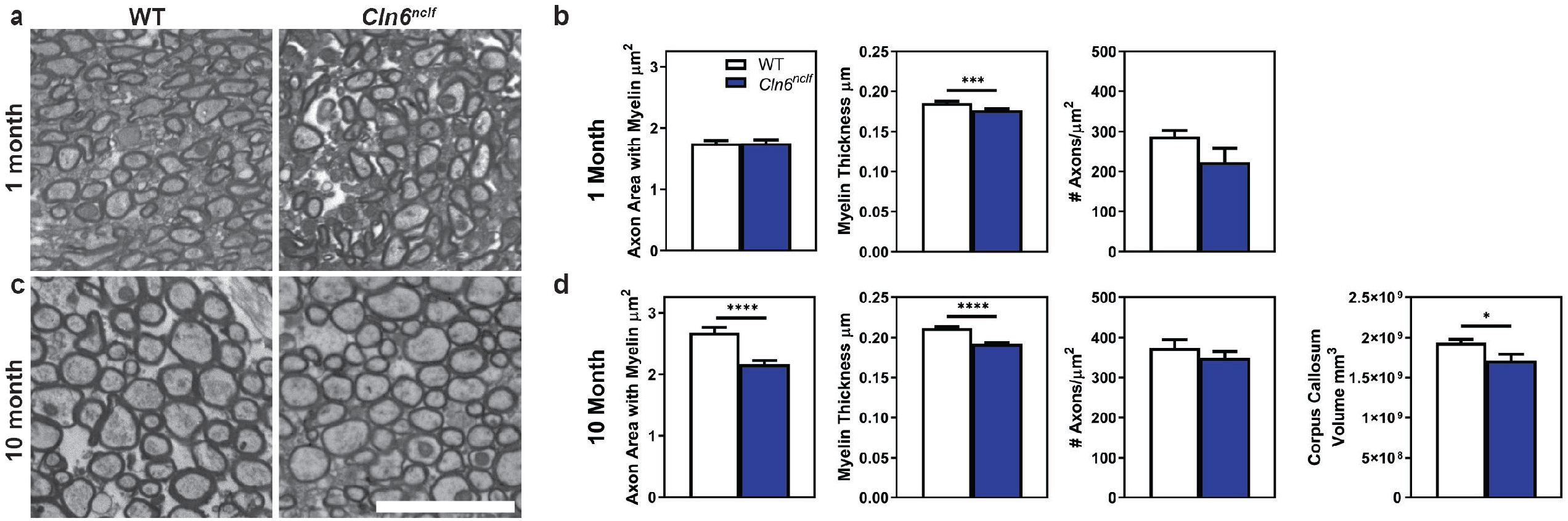
Loss of CLN6 causes myelin deficits in corpus callosum axons. a) Representative images of corpus callosum axonal crossings in one month old wild type and *Cln6*^*nclf*^ brains. b) Loss of CLN6 subtly reduces myelin thickness in the corpus callosum. c) Representative images of corpus callosum axonal crossings in ten month old wild type and *Cln6*^*nclf*^ brains. d) Loss of CLN6 reduces axon and myelin area, myelin thickness, and corpus callosum volume in ten month old animals. Mean ± SEM, unpaired student’s t-test. *p<0.05, ***p<0.001, ****p<0.0001. Scale bar: 5μm

### Lanthionine ketimine ester partially corrects defects caused by CLN6 deficiency

Neuronal polarization, neurite initiation and arborization, and ER-derived vesicle trafficking are all dependent on microtubule transport [35, 37, 38]. Based on this information and our identification of the CCK complex, we hypothesized that the phenotypes we observed in CLN6 deficient neurons were due to defects in microtubule transport resulting from altered CRMP2 function. Thus, we explored whether a CRMP2 modulator could restore some of the altered CRMP2 interactions in CLN6 deficient cells, thereby restoring ER-derived vesicle trafficking in axons and neurite morphogenesis. Additionally, we tested whether CRMP2 modulation could act as a therapeutic target in Batten disease associated phenotypes *in vivo*.

Lanthionine ketimine ester (LKE) binds CRMP2 directly to modulate CRMP2 function and has been found to have therapeutic utility in several neurodegenerative disease models, including Alzheimer’s disease and Multiple Sclerosis [39-43]. LKE has also been shown to partially rescue many of the phenotypes that result from substitution of a hypomorphic CRMP2 allele in *C. elegans*, suggesting that the compound has the potential to restore CRMP2 function in certain conditions [44]. While the use of LKE to correct defects in axonal transport has not been tested, we hypothesized that LKE would restore many of the CRMP2 protein interactions that appear to be disturbed in the absence of CLN6.

When we applied LKE (100 µM) to cultured *Cln6*^*nclf*^ cortical neurons, the number of ER-derived vesicles moving along the axon was restored to wild type levels, as were the levels of ER-tracker signal in the neuronal soma (**Figure 6a-f)**. However, LKE was unable to rescue the levels of PPT1 lysosomal enzyme activity in the synaptic fraction, indicating remaining deficits in the transport of lysosomal cargo (**Figure 6g**). Axonal morphology, as assessed by Sholl analysis, was partially rescued by LKE administration (axon length), however, LKE did not rescue other axonal deficits in *Cln6*^*nclf*^ cortical neurons, specifically the number of axon nodes and ends (**Figure 6h-n**).

**Figure 6.**
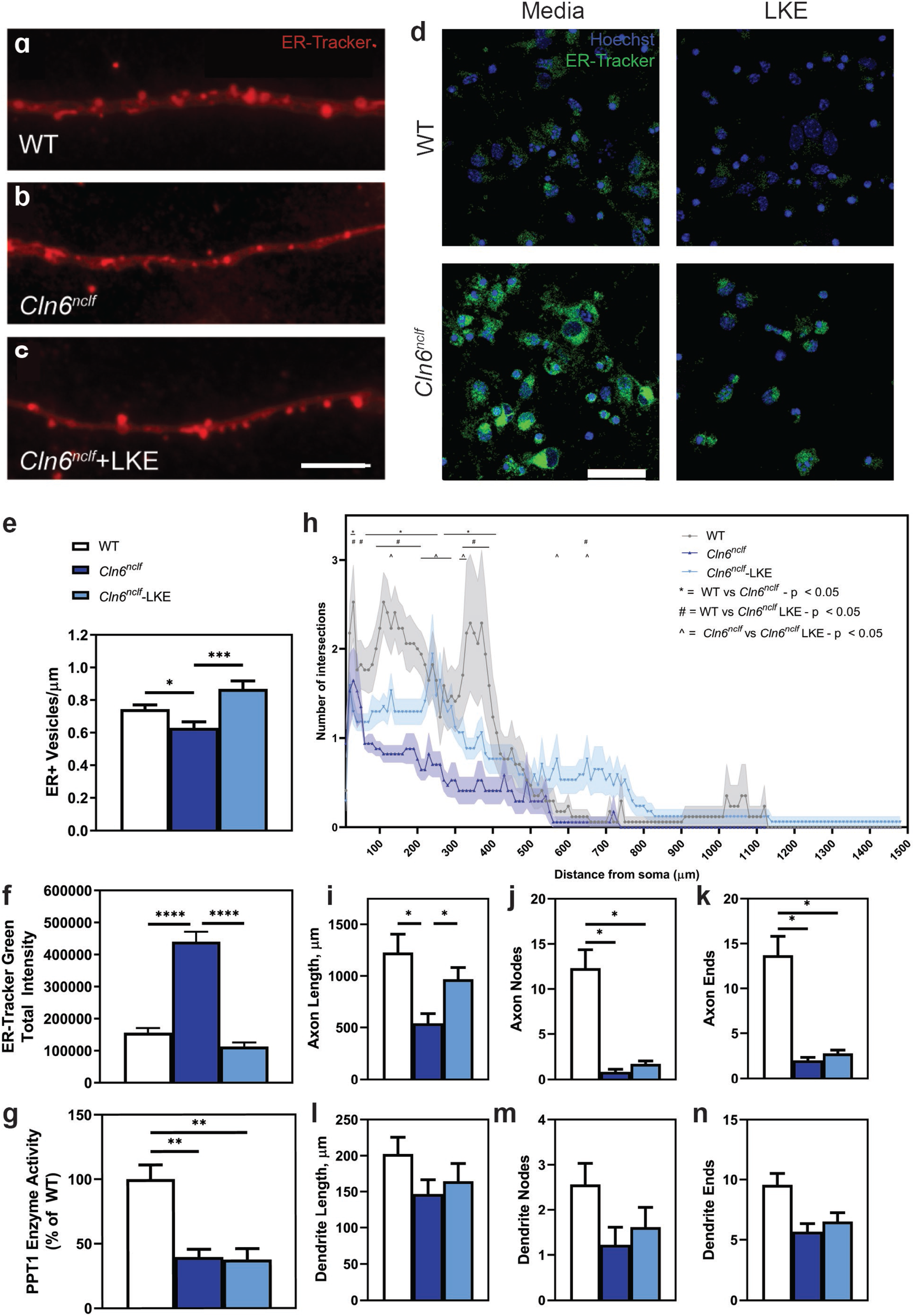
Lanthionine ketimine ester (LKE) restores several *Cln6*^*nclf*^ neurite deficits. a-g) Administration of LKE at 100μM restores ER vesicular trafficking deficits in DIV7 *Cln6*^*nclf*^ cortical neurons, in both the axon (a-c, e) and soma (d, f). h) LKE administration partially correct neurite arborization deficits in *Cln6*^*nclf*^ cortical neurons, as measured by Sholl analysis. i-n) Administration of LKE restores axon length, but not the number of axon nodes and ends in *Cln6*^*nclf*^ mice. Mean ± SEM, Two-way ANOVA with Bonferroni’s post hoc test for Sholl analysis, ordinary one-way ANOVA with Tukey post-hoc for all other graphs. *p<0.05, **p<0.01, ***p<0.001, ****p<0.0001. Scale bar a-c: 5μm; d: 50μm

We surmised that restoration of these neuronal phenotypes may be sufficient to rescue some of the behavioral and pathological phenotypes observed in *Cln6*^*nclf*^ mice. We treated cohorts of wild type and *Cln6*^*nclf*^ mice with LKE dosed in chow (150mg/kg/day) beginning at wean to examine *in vivo* effects when delivered after early postnatal development. In a radial arm maze of memory and learning, LKE treatment reduced the number of errors committed by six-month-old *Cln6*^*nclf*^ mice, though treatment did not affect the time or speed it took for the animals to choose the correct arm (**Figure 7a**). LKE treatment also restored visual acuity during a visual cliff test at eight months of age (**Figure 7b**). However, LKE treatment had no effect on motor ability in an accelerating rotarod test (**Figure 7c**), nor did it prolong *Cln6*^*nclf*^ survival, as treated animals ultimately succumbed at the same time as their untreated counterparts (**Figure 7d**).

**Figure 7.**
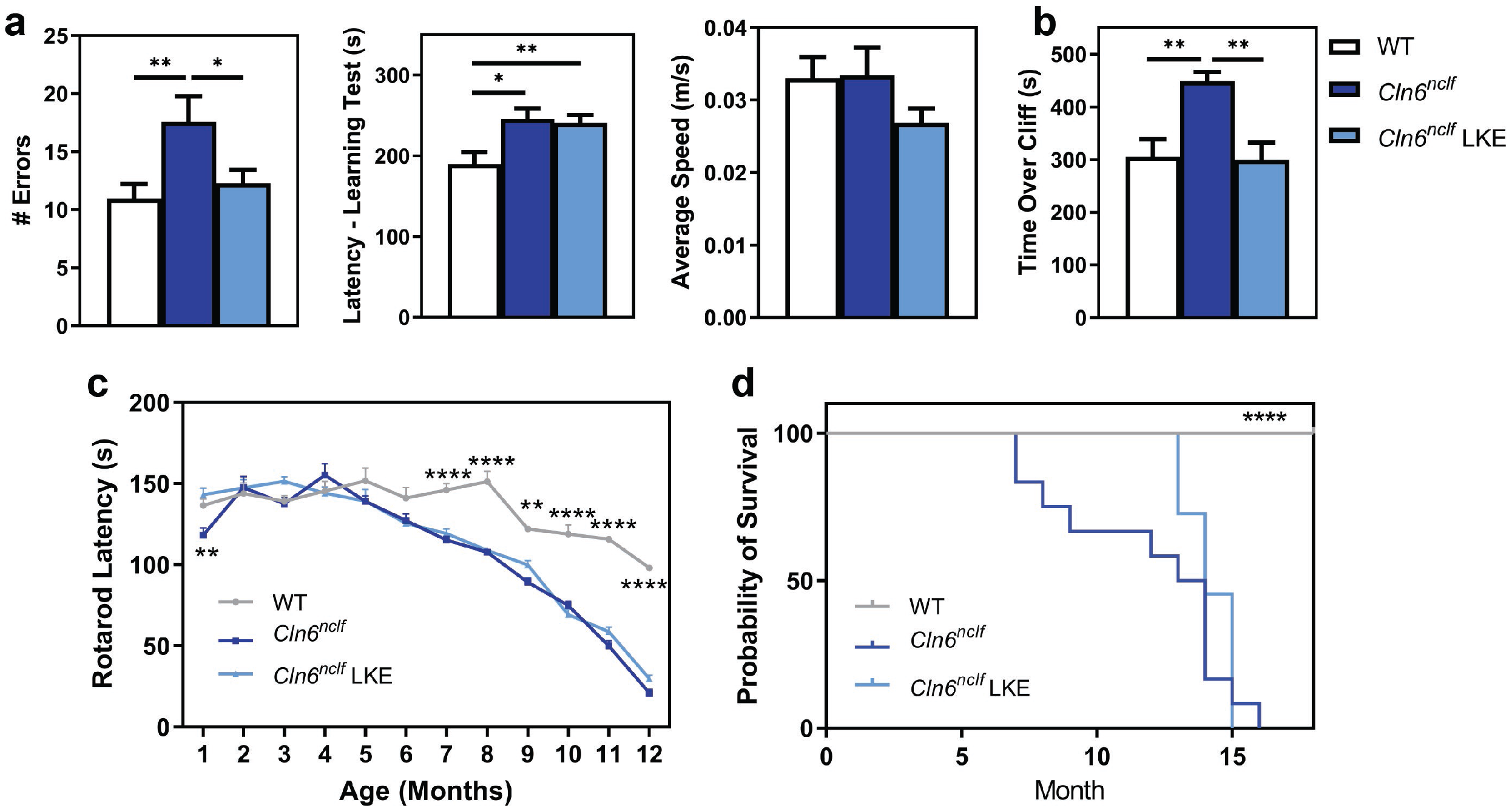
LKE restores some behavioral aspects of *Cln6*^*nclf*^ animals, but ultimately does not extend survival. a) LKE (150mg/kg/day in chow) restores the number of errors made by *Cln6*^*nclf*^ mice in the radial arm maze but does not reduce their latency to choose the correct arm. b) LKE restores *Cln6*^*nclf*^ visual deficits in the visual cliff test. c) LKE does not restore motor deficits in *Cln6*^*nclf*^ mice, as measured by an accelerating rotaroad. d) LKE ultimately does not extend survival of *Cln6*^*nclf*^ mice. Mean ± SEM for all graphs. Ordinary one-way ANOVA with Tukey post-hoc for graphs a-c. Two-way ANOVA with Bonferroni multiple comparisons test was used for graph d. For survival analysis, log rank (Mantel-Cox) test was used. *p<0.05, **p<0.01, ****p<0.0001.

To better understand the etiology behind the behavioral and survival outcomes, histopathology was performed on 11-month-old end-of-life brain samples, as terminal *Cln6*^*nclf*^ mice present with pronounced cortical atrophy, accumulation of ATP synthase subunit C (SubC) in lysosomes, and increased astrocytic and microglial activation at that age [45]. While LKE treatment rescued cortical thickness to wild type levels (**Figure 8a**), microglial activation (CD68^+^), astrocyte reactivity (GFAP^+^), and SubC accumulation were all greatly exacerbated with LKE treatment (**Figure 8b-d**), indicating chronic inflammation and sustained pathology in treated animals.

**Figure 8.**
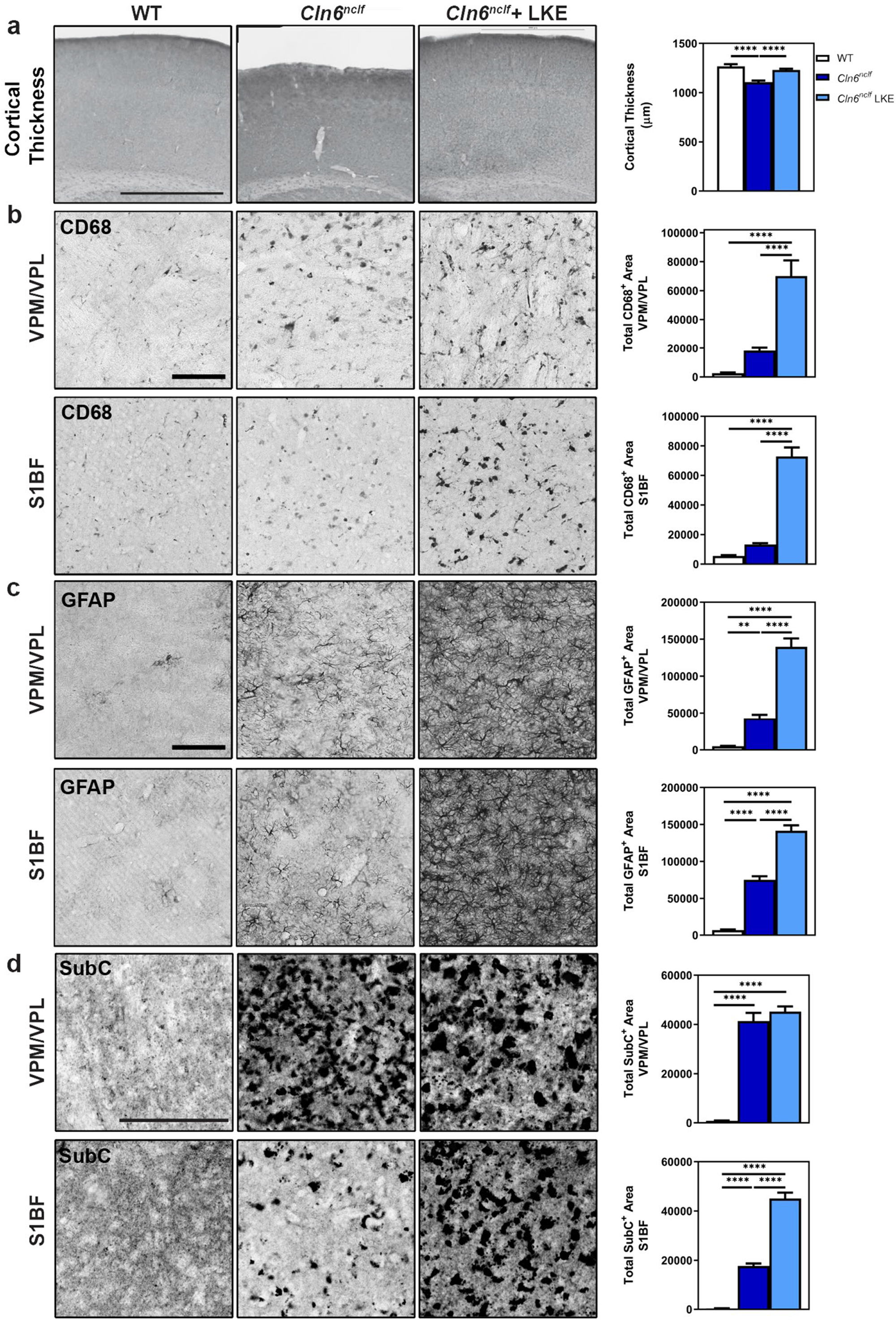
LKE restores cortical thickness in *Cln6*^*nclf*^ mice but does not rescue classic Batten disease pathological measures. a) LKE restores cortical thickness deficits in *Cln6*^*nclf*^ mice. Scale bar: 1000μm. b) LKE enhances microglial response in *Cln6*^*nclf*^ mice in the VPM/VPL of the thalamus and S1BF of the somatosensory cortex. c) LKE enhances astrocyte reactivity in *Cln6*^*nclf*^ mice. d) LKE enhances mitochondrial ATP synthase subunit C accumulation in *Cln6*^*nclf*^ mice. Mean ± SEM, ordinary one-way ANOVA with Tukey post-hoc for all graphs. **p<0.01, ****p<0.0001. Scale bar a: 1000μm; b-d: 100μm

## DISCUSSION

The work presented here demonstrates that CLN6 is a critical regulator of axonal entry of ER-derived vesicles in cortical neurons, likely through interactions with key mediators of vesicle trafficking such as KLC4 and CRMP2.. In the absence of CLN6, dysfunction in this pathway compromises neuronal polarization, neurite extension, and neurite branching. This represents a new etiology in CLN6 disease, and suggests that lysosomal dysfunction is only one component of this complex neurocentric disease. We show that treatment with the CRMP2 modulator LKE is sufficient to rescue some phenotypes associated with CLN6 deficiency in *Cln6*^*nclf*^ mice, supporting the importance of the CLN6-CRMP2 interaction in a neurodegenerative disease and introducing a potential new therapeutic target for CLN6-Batten disease and other disorders of neuronal vesicle trafficking.

While we have established that CLN6 interacts with both CRMP2 and KLC4, the precise nature of these interactions remains to be characterized. CRMP2-KLC4 interactions are markedly increased in the absence of CLN6; thus, one likely possibility is that CLN6 negatively regulates the coupling of CRMP2 to KLC4 and the microtubule network. When CLN6 is absent, CRMP2 may bind more tightly to motor proteins and thus fail to disengage at appropriate times. A similar phenomenon may exist for other CRMP2 interactions, with CLN6 differentially regulating the affinity of CRMP2 for various partners. Further biochemical studies may be able to elucidate these mechanisms at the molecular level.

We observed that CLN6 regulates the axonal abundance of ER-derived vesicles, but not lysosomes, suggesting some degree of selectivity with regard to pools of membrane-bound vesicles. The necessity for CLN6 in neurite polarization, extension, and branching suggests that ER-derived vesicles may be carrying building blocks that are critical for neurite structure. It will be important to determine if the CCK complex regulates the trafficking of a highly diverse, heterogenous ER-derived vesicle pool, or, alternatively, highly specific cargoes. In the latter case, CLN6 could be acting as a compartmental barcode, specifying cargoes for certain destinations.

CLN6 may play dual roles in neurons which could explain why loss of CLN6 predominately presents as a neurodegenerative disorder. In non-neuronal cells, CLN6 works with CLN8 to sort and transport proteins targeted to the lysosome, and loss of either protein results in lysosomal dysfunction [46]. Here, we show in neurons that CLN6 is also required for facilitating transport of ER-derived vesicles from the neuronal soma to the microtubule transport network within axons. While modulation of CRMP2 though LKE successfully increased ER-derived vesicular transport, the lack of functional CLN6 would still be expected to hinder the loading of necessary lysosomal constituents, and the increased flux of cargo could then result in greater lysosomal burden and the observed increases in storage material. Thus, while LKE was beneficial for some aspects of cellular pathology, it would likely be insufficient as a standalone treatment for CLN6 disease.

It has been argued that the most important cellular pathologies in many different lysosomal storage disorders may be unrelated to lysosomes. While the accumulation of autofluorescent lysosomal storage material is a salient hallmark of these disorders, several of these conditions (including CLN6-Batten disease) are caused by mutations in genes encoding proteins that localize to other cellular compartments. In these cases, lysosomal dysfunction may be just one downstream symptom of a more basic cellular etiology. We have shown that CLN6 deficiency interrupts ER-vesicle trafficking, which is important for a host of cellular processes. One consequence of this dysfunction may be that critical cargoes are not delivered to the lysosomal compartment, but it is important not to ignore the myriad of other processes that are also dependent on vesicle-mediated trafficking. While working towards the goal of developing therapies which correct the cellular pathologies in various forms of lysosomal storage disorders, identifying and targeting affected pathways that are upstream of the lysosome could prove to be a powerful strategy.

## Acknowledgements

This work was supported through the National Institutes of Health (R01NS082283, P20GM103620, and P20GM103548.

## MATERIALS & METHODS

### Animals

Animal protocols were approved by the Sanford Research Institutional Animal Care and Use Committee (USDA License 46-R-0009) with all procedures carried out in strict accordance with National Institutes of Health guidelines and the Sanford Research Institutional Animal Care and Use Committee guidelines. Wild type (WT) C57BL/6J and homozygous *Cln6*^*nclf*^ mice on a C57BL/6J background were obtained from the Jackson Laboratory. Lanthionine ketamine ester was provided as a gift by Hensley, K.

### Yeast Two Hybrid

The hydrophilic N-terminal region of CLN6 was screened against a human fetal brain library using Stratagene’s Cytotrap yeast two-hybrid system (Y2H) as previously published (Stratagene, La Jolla, CA). Briefly, this Y2H system utilizes the temperature sensitive yeast strain, Cdc25H, which harbors a mutation in the homolog of the human Sos (hSos) protein, Cdc25p. The N-terminus of CLN6 is fused with hSos and when brought to the plasma membrane through an interaction with myristoylated library protein, restores growth at 37°C.

### Cell Culture

*<u>PC12 Cells:</u>* Rat pheochromocytoma (PC12) cells (ATCC; Manassas, VA) were cultured at 37°C with 5% CO_2_ on plastic culture dishes in DMEM/F12 medium supplemented with 10% heat inactivated horse serum, 5% fetal bovine serum, and penicillin/streptomycin. PC12 cells were differentiated into neuron-like cells on plastic culture dishes coated in rat tail collagen or glass coverslips coated with 50 µg/mL of Poly-L-lysine while being cultured in serum/antibiotic free DMEM/F12 media supplemented with 50 ng/mL nerve growth factor (NGF). *<u>Primary cortical neurons:</u>* Mouse cortical neurons were prepared at mouse embryonic day (E) 15.5 as previously described [47]. Briefly, both cerebral hemispheres were harvested and cerebral cortices were isolated in ice cold complete HBSS containing 2.5mM HEPES, 30mM D-glucose, 1mM CaCl_2_, 1mM MgSO_4_ and 4mM NaHCO_3_ in HBSS. The tissues were digested by papain solution at 37°C, triturated, passed through a 70µm meshed cell strainer, and plated on coverslips precoated with poly-D lysine and Laminin. Neurons were maintained in Neurobasal media supplemented with B27 and 2mM L-glutamine at 37°C in 5% CO_2_. Half of the media was changed on alternating days. *<u>Live cell imaging</u>*. E15.5 primary cortical neurons were plated at a density of 25,000 cells/well or 50,000 cells/well in 35mm glass bottom dishes. Live imaging was performed at 2 and 7 days *in vitro* (DIV). ER-derived vesicles were visualized with ER-Tracker Red dye (Invitrogen Molecular Probes) according to the supplied protocol. Using a 60X oil-immersion objective on a Nikon A1 inverted confocal microscope, images were captured every 2 seconds for 150 seconds. Kymograph analysis was performed with Metamorph (Molecular Devices) to determine vesicle velocity. *Coimmunoprecipitation and Western Blotting*. CoIPs and Western blotting were performed as previously described (48). Briefly, wild type and *Cln6*^*nclf*^ cortices were homogenized in Lysis/IP buffer and protein concentrations were quantified by the Pierce 660 nm assay (Fischer; Rockford, IL) using a SpectraMax M5 plate reader. Synaptosomal preparations were isolated with Syn-PER buffer (Thermo), according to the manufacturer’s protocol, and were homogenized in Lysis/IP buffer as above. Protein A/G PLUS-Agarose beads were cleared of storage buffer and resuspended in diluted antibody of interest for 1 hour at 4°C with rotation. Beads were pelleted and washed with wash buffer, resuspended in Lysis/IP buffer, and incubated with lysate overnight at 4°C with rotation. Beads were then washed, resuspended in 2X sample buffer, and heated to 100°C for 5 minutes, cooled, and briefly centrifuged with the supernatant moved to a clean tube for analysis by Western blot. 10 µg of protein was loaded onto either fixed percentage or gradient SDS-PAGE gels. Gel samples were transferred to nitrocellulose membranes and blocked with TBS-T plus 5% non-fat dry milk followed by overnight incubation in primary antibodies. Samples were washed and incubated with appropriate horseradish peroxidase (HRP) conjugated secondary antibodies. Membranes were then subjected to enhanced chemiluminescent detection (ECL) via ECL Western Blotting Detection Reagents via the manufacturer’s protocol (GE Healthcare, Amersham; Piscataway, NJ). Blots were imaged and captured using the UVP Biospectrum 500 imaging system (Upland, CA), and total pixel density within a defined region was measured and compared against a β-actin control to generate total density ratio. VisionWorks analysis software was used to generate data of total density values with Microsoft Excel used to calculate ratios.

### Antibodies

The following antibodies were used for immunoblotting and immunohistochemistry: anti-CRMP2 1B1 (Abcam ab62539), anti-CRMP2 (Cell Signaling 9393), anti-Phospho-CRMP-2 (T514 (Cell Signaling 9397), S522 (ECM BioSciences CP2191)), anti-Numb mAb (Cell Signaling C29G11), anti-Neurofibromin (NF1) (Millipore 05-1430, clone NFn27), anti-Neural Cell Adhesion Molecule L1 (Millipore MAB5272, clone 324), anti-PIR121/Sra-1 (Upstate 07-531), anti-KLC4 (Abcam ab89040), anti-KLC4 (Santa Cruz sc-134680 H-54), and anti-β-Actin (Cell Signaling 4967).

### Plasmids and Electroporations for Cell Visualization for Sholl Analysis

pCAG-GFP, a gift from Connie Cepko (Addgene plasmid # 11150 [43]), was dissolved in 10mM Tris-HCl (pH 8.0). For *in vitro* Sholl analysis, *ex utero* electroporation was used to deliver DNA as previously described. E15.5 embryos were harvested in cold complete HBSS, and 1μL of 2μg/μL plasmid with 0.1% Fast Green (Sigma) was injected by hand into the lateral ventricle using a Picospritzer II (Parker). Five 50ms pulses (one second apart) of 45V were applied across the two cerebral hemispheres using an Electro Square Porator ECM 830 (BTX) fitted with a tweezer-type electrode. Primary cortical culture were processed as described. *In utero* electroporations were performed similarly, according to a previously reported protocol [44].

### Immunohistochemistry and Immunocytochemistry

IHC and ICC experiments were performed as previously described [45, 48]. Briefly, for the ICC experiments, adherent neurons were fixed with 2% paraformaldehyde (PFA), 2% sucrose in 0.1M PBS for 20 minutes on ice, and rinsed. For *in vivo* studies, animals were CO_2_ euthanized and transcardially perfused with 4% paraformaldehyde and brains post-fixed at 4°C for 24 hours. Brains were sliced at 50 µm thickness on a vibratome (Leica), coverslipped, and blocked for one hour as previously described. Samples were incubated in primary antibodies overnight at 4°C, and included: anti-ATP5G1 (Fisher PA5-25799), anti-GFAP (Dako, M0761), and anti-CD68 (AbD Serotec, MCA1957). Cell dyes included ER tracker-Red (Invitrogen Molecular Probes) and or ActinGreen (Invitrogen Molecular Probes). Samples were then washed and incubated with 4,6-diamidino-2-phenylindole (DAPI, Thermo Fisher)) and Alexa-fluor conjugated secondary antibodies (Invitrogen, Thermo Fisher) for 1 hour at room temperature. Samples were washed and mounted for imaging with Dako Faramount aqueous mounting medium (Agilent Technologies, Lexington, MA). Tissue sections were imaged using a Nikon Eclipse Ni-E upright microscope using a 20x dry objective lens, and cultured neurons were imaged using a Nikon A1 inverted confocal microscope using 60x and 100x oil immersion lenses, with the pinhole set to 1 airy unit (AU).

### Image Quantification

For quantification of neurite arborization, neurons were traced using Neurolucida software (MBF Bioscience, Williston, VT) and Scholl analysis was conducted using Neurolucida Explorer software as previously described [49]. Cortical thickness was measured in triplicates from the pial surface through layer 6 of the cerebral cortex. SubC, GFAP, and CD68, tissues were imaged and analyzed using a Nikon 90i microscope with NIS-Elements Advanced Research software or equivalent, and threshold analysis performed.

### Electron Microscopy

Volume of the Corpus Callosum was estimated using the Calavieri Estimator and Stereoinvestigator[50]. In brief, PBS-perfused mouse brains were split along the midline into hemispheres and one hemisphere was fixed in 4% PFA. 50 micron sagittal sections, originating from the midline, were sectioned and stained with Hematoxylin and Eosin. For the Calavieri estimation every fifth section was used with a standard grid size. The Gundersen coefficient of error (M=1) was less than 10% for each hemisphere sample (n = 3 mice per group).

### Behavioral Tests

The radial arm maze and visual cliff were captured using Any-Maze software, while the accelerating rotarod was manually recorded by a blinded experimenter. *<u>Radial Arm Maze:</u>* Radial arm was performed as previously described, where animals were trained to remember where three sucrose pellets were placed from previous trials [51]. The number of errors (ie, number of entries into an arm without a sucrose pellet), the latency to complete the task, and speed were recorded. *Visual Cliff:* Visual Cliff was performed as previously described [30]. Briefly, mice were placed in a constructed wooden box with plexiglass bottom, balanced over the ledge of a table. Mice were observed for the amount of time spent over the ‘visual cliff’ of the table ledge in a single, 15 minute trial. *Rotarod:* Rotarod motor testing was performed as previously described, with an acceleration to 30rpm over a period of 240 seconds [30]. The latency to fall from the rotating rod was recorded over the course of nine trials.

### Statistical Analyses

At least three animals were analyzed for each genotype. All quantifications are reported as the mean ± the standard error in the mean (SEM). Statistical analysis was performed in Graphpad Prism 7 or later. Student’s t-tests were performed as two-tailed tests, with F-tests used to compare variances and p<0.05 used as the cutoff for significance. One-way ANOVAs were performed, with Tukey’s multiple comparison post-hoc correction comparing all possible groups. Two-way ANOVAs were performed, with Bonferroni’s multiple comparison post-hoc test. *p<0.05, **p<0.01, ***p<0.001, ****p<0.0001.

## FIGURE LEGENDS

**Supplemental Figure 1 – Loss of CLN6 does not alter pCRMP2 dynamics**. a) Loss of CLN6 does not alter total CRMP2 or CRMP2 phosphorylation at multiple residues. b) Quantification of a. Mean ± SEM, unpaired student’s t-test.

