## Supplementary figures and images for "A CLN6-CRMP2-KLC4 complex regulates anterograde ER-derived vesicle trafficking in cortical neurites"

### Supplemental Figure 1

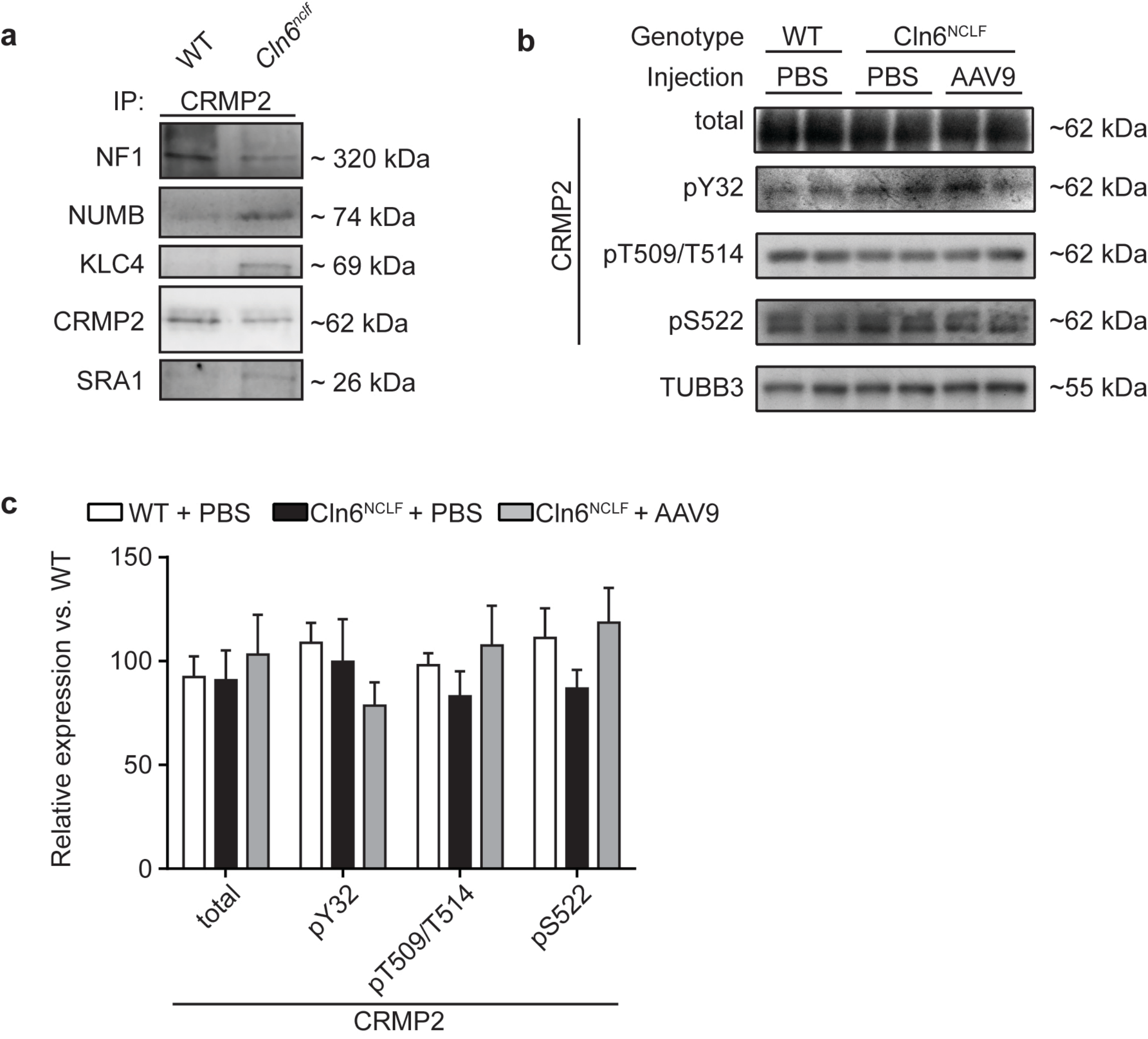
